# Disrupting *Pitx1* regulatory topology results in overtly normal limb development

**DOI:** 10.1101/138644

**Authors:** Richard Sarro, Deena Emera, Severin Uebbing, Emily V. Dutrow, Scott D. Weatherbee, Timothy Nottoli, James P. Noonan

## Abstract

Gene expression patterns during development are orchestrated in part by thousands of distant-acting transcriptional enhancers. However, identifying enhancers that are essential for expression of their target genes has proven challenging. Genetic perturbation of individual enhancers in some cases results in profound molecular and developmental phenotypes, but in mild or no phenotypes in others. Topological maps of long-range regulatory interactions may provide the means to identify enhancers critical for developmental gene expression. Here, we leveraged chromatin topology to characterize and disrupt the major promoter-enhancer interaction for *Pitx1*, which is essential for hindlimb development. We found that *Pitx1* primarily interacts with a single distal enhancer in the hindlimb. Using genome editing, we deleted this enhancer in the mouse. Although loss of the enhancer completely disrupts the predominant topological interaction in the *Pitx1* locus, *Pitx1* expression in the hindlimb is only reduced by ~14%, with no apparent changes in spatial distribution or evidence of regulatory compensation. *Pitx1* enhancer null mice did not exhibit any of the characteristic morphological defects of the *Pitx1*^−/−^ mutant. Our results indicate that *Pitx1* expression is robust to the loss of its primary enhancer interaction, suggesting disruptions of regulatory topology at essential developmental genes may have mild phenotypic effects.

## Introduction

Embryonic development depends on the spatial, temporal, and quantitative control of gene expression. The complex regulatory programs that orchestrate morphogenesis are specified by thousands of tissue-specific, distant-acting enhancers (1–4), which are brought into contact with their gene targets via long-range chromatin looping interactions (3–12). These contacts occur within stable, megabase-scale topologically associated domains (TADs), many of which are conserved across tissues and species (13–16). Despite their clear overall importance for development, quantifying the contribution of individual enhancers to the expression of their target genes has been difficult. Genetic knockout studies of single enhancers in the mouse have yielded a range of molecular and phenotypic effects. In a classic example, deletion of the ZRS, a long-range enhancer that controls expression of *Shh* specifically in the developing limb bud, results in loss of *Shh* expression and truncation of the mouse limb (17, 18). However, knockouts of other enhancers near developmental genes show less severe phenotypes (19, 20) or no obvious phenotype at all (21).

One potential mechanism to account for this result is that many developmental genes are regulated by multiple, redundant enhancers with overlapping functions that may compensate for the loss of a single enhancer (5, 22–26). This redundancy potentially serves to buffer the effects of genetic disruptions of individual enhancers. Clearly, however, the loss of some enhancers cannot be buffered; the ZRS appears to be the primary source of regulatory information for specifying *Shh* expression in the limb bud (17, 18). Identifying such critical enhancers that are likely to exhibit large-effect loss of function mutations has been challenging. To date there is as yet no means to distinguish enhancer mutations with large effects from mutations that can be buffered by compensatory mechanisms. This presents a major barrier for efforts to understand the contribution of regulatory variation to human disease.

We hypothesized that topological interactions would be a useful filter, in that individual enhancers critical to tissue-specific expression of pleiotropic developmental genes would show robust, tissue-specific interactions with those targets. In this way topology may be used to identify not just an enhancer’s targets, but also to predict the potential effects of an enhancer deletion. Supporting this hypothesis, it has been shown that the ZRS directly interacts with *Shh* in the limb bud (27, 28) and *HoxD* gene expression in the developing limb is specified by an array of interacting enhancers (5). To identify potentially essential developmental enhancers, we leveraged the results of a recent study that utilized Chromatin Interaction Analysis by Paired-End Tag Sequencing (ChIA-PET) (9, 29) targeting the SMC1a cohesin subunit to identify over 2,500 cohesin bound chromatin loops in E11.5 mouse embryonic limb (10). From this study we identified a robust, hindlimb-specific interaction between the pleiotropic developmental gene *Pitx1* and a distal sequence, which we termed the *Pitx1* Distal Enhancer (PDE), located 132 kb away.

*Pitx1* encodes a homeodomain transcription factor expressed in the hindlimb and mandibular arch (30, 31). *Pitx1*^*+/−*^ mice are overtly normal, exhibiting only minor morphological defects at a low levels of penetrance (32). *Pitx1*^*−/−*^ mice show early postnatal lethality, as well as loss of the ilium, a reduction in length of the femur, a loss of the patella, and a reduced, malformed mandible (30, 31). *Pitx1* is hypothesized to establish hindlimb identity, and overexpression of *Pitx1* in the forelimb produces a hindlimb-like morphology (33). Prior studies have identified *Pitx1* enhancer mutations that result in substantial morphological phenotypes in non-mammalian systems (34, 35). We predicted loss of the PDE, which our results indicate is the major topological interaction partner of the *Pitx1* promoter in the hindlimb, may also have large molecular and morphological effects. The PDE displays strong, hindlimb-specific H3K27ac marking, a signature of active enhancers; in the forelimb the sequence is marked by H3K27me3, associated with PRC2-mediated repression (36). The PDE is also highly conserved across amniote species, suggesting it serves an ancient role in *Pitx1* regulation.

We used CRISPR to delete the PDE in the mouse, thus abolishing any regulatory input or topological insulation conferred by the interaction. We anticipated two potential outcomes from the loss of the PDE: the recapitulation of the morphological defects of the gene deletion, or the formation of aberrant and/or compensatory interactions between *Pitx1* and other potential regulatory elements in the absence of the primary interaction. Instead, we found the loss of the PDE to have mild effects on *Pitx1* expression in the hindlimb and insignificant effects on hindlimb morphology. We did not identify changes in enhancer activity or new long-range interactions, at least at a level detectable by our methods, that may have compensated for the loss of the PDE.

## Results

### *Mapping* Pitx1 *regulatory interactions*

We first characterized the topological landscape of the *Pitx1* promoter in the mouse embryonic forelimb and hindlimb using circularized chromosome conformation capture (4C) (37). We were able to recapitulate the previously described *Pitx1*-PDE interaction in the wild type mouse E11.5 hindlimb. This interaction is the most prominent topological interaction we identified involving the *Pitx1* promoter (FourCseq replicate 1 *P* = 0.00754, replicate 2 *P* = 0.00374; see Materials and Methods; Fig. 1 and Table S1). *Pitx1* also interacts with the PDE at a low level in the E11.5 forelimb (FourCseq replicate 1 *P* = 0.03067, replicate 2 *P* = 0.05542). These interaction frequencies correlate with the expression levels of *Pitx1* and epigenetic profile of the enhancer in the hindlimb and forelimb. In the hindlimb, *Pitx1* is expressed and the PDE exhibits high levels of H3K27ac (Fig. 1). Conversely, *Pitx1* is not expressed in the forelimb, and both the *Pitx1* promoter and the PDE sequence are marked by H3K27me3, indicating they are repressed in this tissue (36).

**Figure 1.**
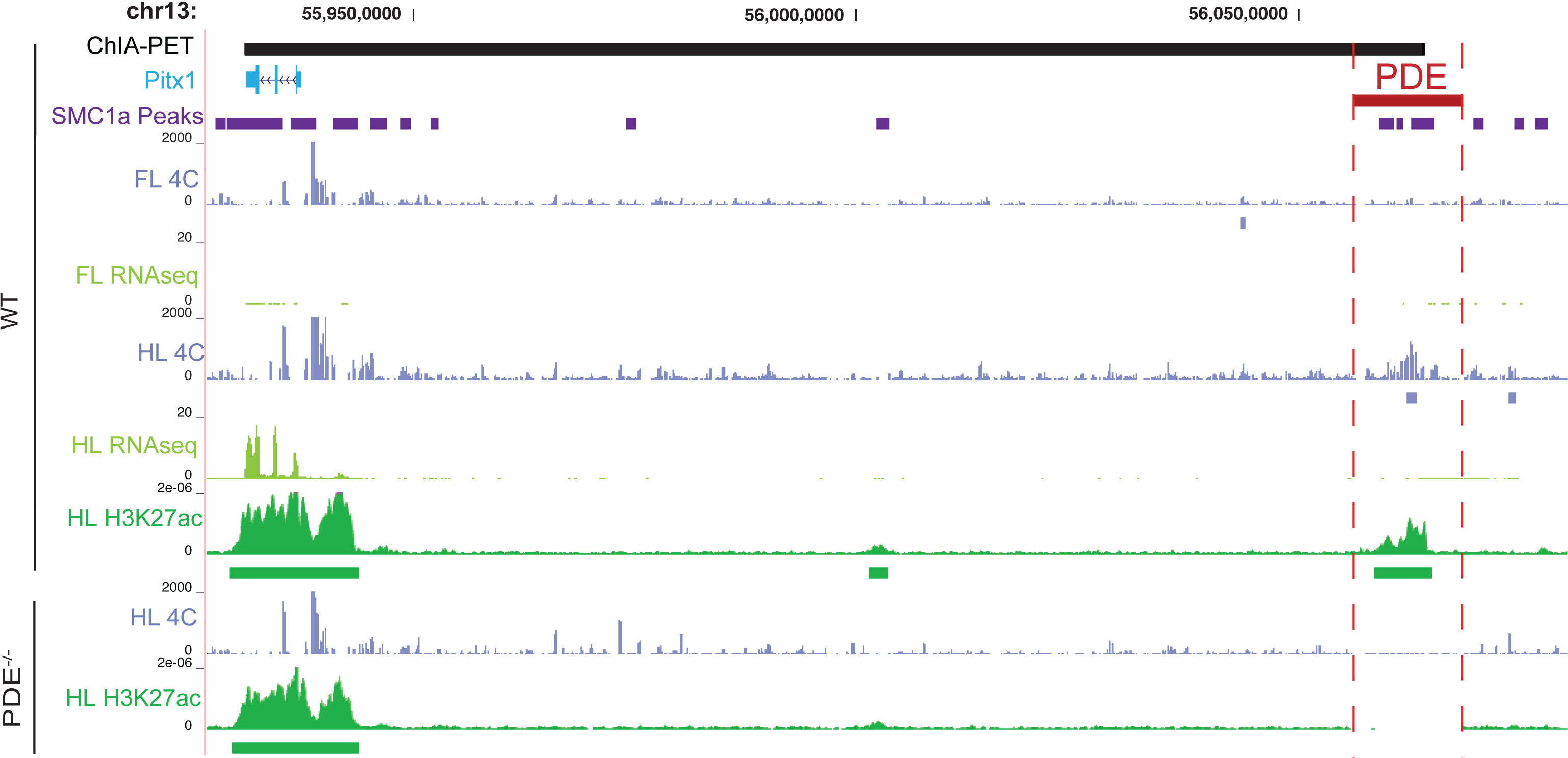
The *Pitx1* regulatory landscape in wild type and PDE−/− mouse E11.5 limb buds. *Top*. 4C interaction profiles of the *Pitx1* promoter in wild type forelimb and hindlimb are shown in blue. Reproducible, nominally significant interacting regions as called by FourCSeq are shown below each track. Transcription profiles (RNA-seq) in forelimb and hindlimb are shown in light green, and H3K27ac signal and enriched regions in hindlimb are shown in dark green. The ChIA-PET interaction between *Pitx1* and the PDE previously identified in E11.5 limb buds is shown in black at the top of the figure, and SMC1a binding sites are shown in purple (10). The location of the PDE (chr13:56,055,928-56,068,947 in mm9) is shown in red. *Bottom. Pitx1* promoter interaction profile and H3K27ac signal in PDE−/− E11.5 hindlimb.

### *Generation and validation of* PDE^−/−^ *mice*

In order to disrupt the *Pitx1*-PDE interaction, we used CRISPR/Cas9 genome editing to generate a Cre-loxP conditional deletion allele in C57BL6/J mice (Fig. S1). The locations of the loxP insertions were designed to ensure that all local SMC1a binding sites identified would be removed (10). We first validated the deletion of the PDE sequence via PCR and Sanger sequencing (Fig. S1). We then validated the ablation of the topological interaction using 4C (Fig. 1). The loss of the PDE sequence is also evident as a lack of sequencing reads in the deleted interval in limb H3K27ac ChIP-seq and 4C datasets (Fig. S1, Fig. 1). We did not detect any evidence of increased interaction frequency in the genomic regions flanking the deletion, supporting that removal of the PDE completely disrupts the promoter-enhancer interaction.

Although *Pitx1*^*−/−*^ mice exhibit neonatal lethality, we did not observe any reduction in viability in constitutive PDE^−/−^ mice. PDE^+/−^ × PDE^+/−^ crosses yielded wild type, PDE^+/−^ and PDE^−/−^ offspring at the expected 1:2:1 ratio (Table S2). We therefore compared wild type and constitutive homozygous knockout mice in all subsequent analyses.

### *Comparison of* Pitx1 *expression in wildtype and PDE*^−/−^ *mice*

We next considered the effects of the PDE deletion on *Pitx1* expression. We examined two tissues that are known to both express *Pitx1* during embryonic development and exhibit malformations in *Pitx1*^*−/−*^ mice: the hindlimb and the mandibular arch. To analyze *Pitx1* expression, we initially compared hindlimbs and mandibular arches from litter-matched wild type and PDE^−/−^ mice using RT-qPCR (Fig. 2, Fig. S2). We carried out a time course analysis of *Pitx1* expression in the hindlimb from E10.5 through E13.5. *Pitx1* expression was reduced by 13-16% in PDE^−/−^ hindlimb at all four time points. This reduction was a significant effect of the PDE deletion, and not solely attributable to litter effects or variation across developmental time points (three-way mixed effect ANCOVA F_1,20_=18.624, *P* = 0.000336; Fig. 2; see Materials and Methods). We also carried out pairwise comparisons at each time point, which indicated that *Pitx1* expression was significantly reduced in PDE^−/−^ hindlimb at E12.5 and E13.5 (Mann-Whitney U test *P* = 0.012 and *P* = 0.021, respectively; Fig. S3) (38). Loss of *Pitx1* expression was more substantial in PDE^−/−^ E11.5 mandibular arch (39% expression reduction, Mann-Whitney U test *P* = 0.00086).

**Figure 2.**
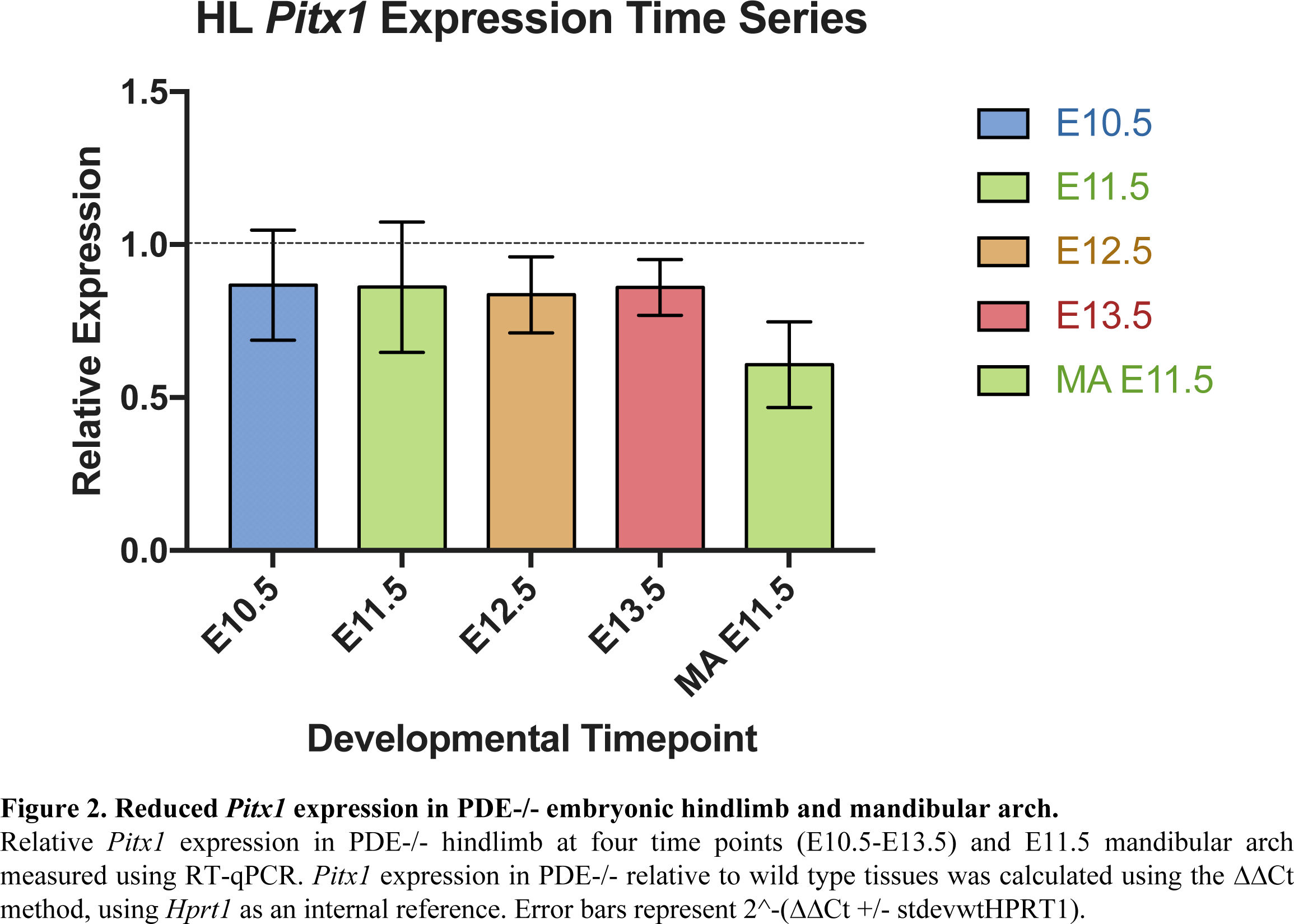
Reduced *Pitx1* expression in PDE−/− embryonic hindlimb and mandibular arch. Relative *Pitx1* expression in PDE−/− hindlimb at four time points (E10.5-E13.5) and E11.5 mandibular arch measured using RT-qPCR. *Pitx1* expression in PDE−/− relative to wild type tissues was calculated using the ΔΔCt method, using *Hprt1* as an internal reference. Error bars represent 2^-(ΔΔCt +/− stdevwtHPRT1).

We then performed RNA-seq in wild type and PDE^−/−^ hindlimb and mandibular arch at E11.5 to identify potential global changes in gene expression due to reduced *Pitx1* levels. In PDE^−/−^ mice, *Pitx1* showed a similar reduction in expression in both tissues as we observed by RT-qPCR (14.5% in hindlimb, 32.14% in mandibular arch; Table S3-S4). However, the change of expression in each tissue was not statistically significant after multiple testing correction (hindlimb DESeq Benjamini-Hochberg corrected (BH) *P* = 0.9997, mandibular arch DESeq BH *P* =0.5946, (39)). Expression of *Tbx4*, a known downstream regulatory target of *Pitx1* in the hindlimb (33), was reduced by 4.68% (p=0.295, BH *P* = 0.9997). Our DESeq analysis did not identify any genes whose expression was significantly altered between the wild type and PDE^−/−^ after multiple testing correction in either tissue (BH *P* < 0.05.).

To gain insight into potential subtle effects of *Pitx1* reduction of expression, we conducted Gene Ontology (GO) enrichment analyses of all genes exhibiting expression changes with a non-adjusted *P* value < 0.05 (271 hindlimb genes, 234 mandibular arch genes; Table S5). Genes that met this relaxed threshold in either tissue were associated with GO categories related to embryonic development and gene regulation. However, no ontologies reached a BH adjusted *P* value < 1.5E-07 (40). This further suggests the reduced dosage of *Pitx1* did not cause substantial changes to regulatory networks or developmental processes in either the limb or the mandibular arch.

To determine if loss of the *Pitx1*-PDE interaction resulted in changes in the spatial distribution of *Pitx1* expression in the hindlimb and mandibular arch, we visualized *Pitx1* expression using whole mount *in situ* hybridization on litter-matched wild type and PDE^−/−^ E11.5 embryos. In agreement with previous studies (30), our *in situ* analysis detected *Pitx1* expression localized to the mandibular arch and hindlimb in wild type embryos (Fig. 3). We observed the same pattern of expression in PDE^−/−^ embryos, indicating that the spatial distribution of *Pitx1* expression at E11.5 is not substantially altered due to loss of the PDE (Fig. 3, Fig S4).

**Figure 3.**
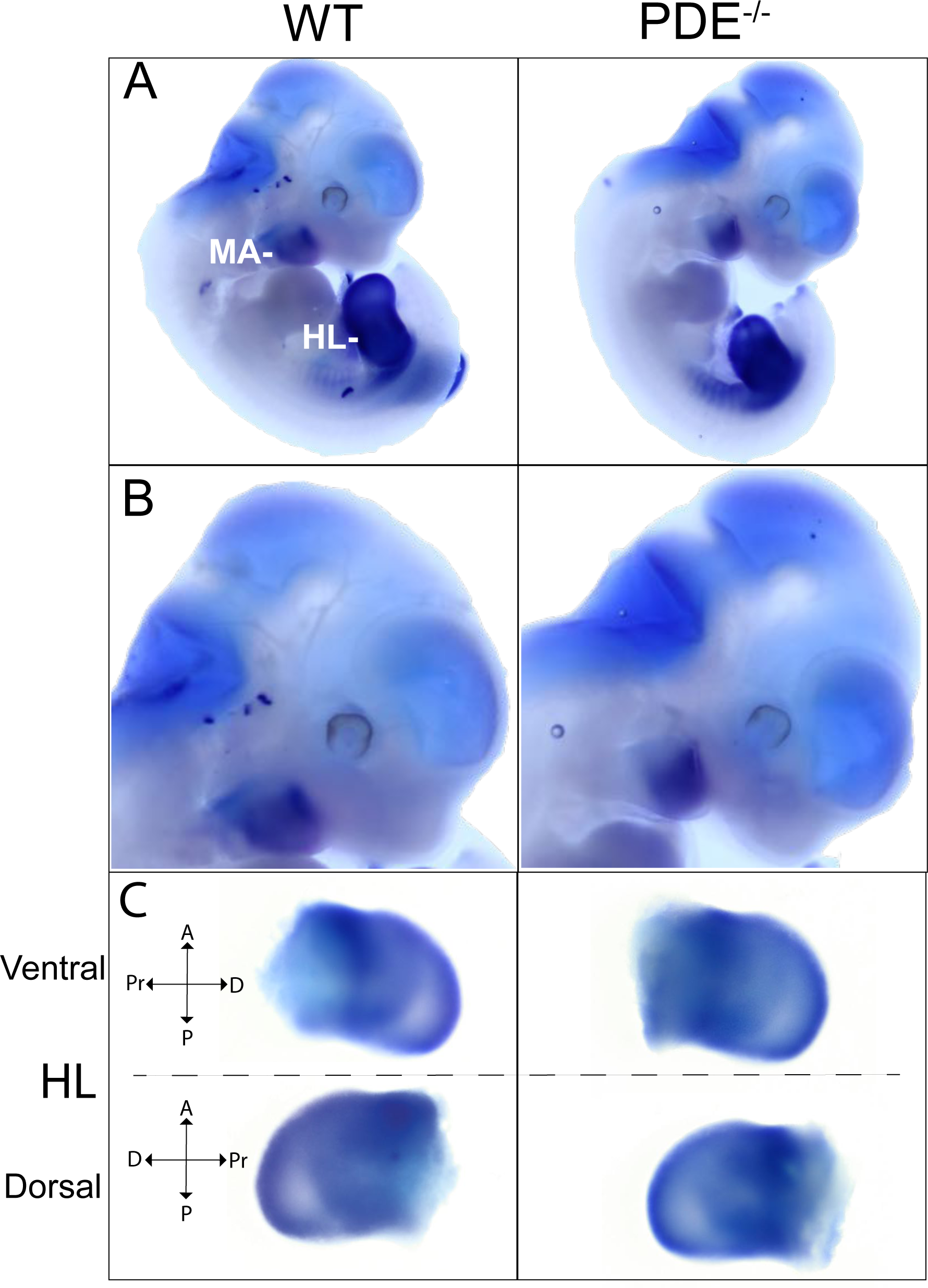
Spatial distribution of *Pitx1* in wild type and PDE−/− E11.5 embryos. A. *Pitx1* expression in wild type and PDE−/− embryos visualized using whole mount *in situ* hybridization. The hindlimb (HL) and mandibular arch (MA) expression domains are labeled. B. Magnified view of *Pitx1* expression in the mandibular arch in the embryos shown in A. C. *Pitx1* expression in wild type and PDE−/− E11.5 hindlimb. Ventral and dorsal views are shown. A = anterior; P = posterior; Pr = proximal; D = distal. Background surrounding each embryo in each photograph was removed for purposes of visualization.

### Comparing epigenetic signatures of regulatory element activity between WT and PDE^−/−^ mice

Deletion of the PDE sequence removed the major interacting partner of the *Pitx1* promoter we could detect in the hindlimb. We hypothesized this loss would result in an unoccupied chromatin anchor point at the *Pitx1* promoter, leaving it potentially accessible to other regulatory interactions that may compensate for loss of the primary enhancer. Such compensatory events might be detected as changes in the observed interaction frequency and epigenetic profile at other loci within the TAD containing *Pitx1* (13). We did not observe any robust novel topological interactions in PDE^−/−^ mice at a level detectable by 4C (FourCSeq P < 0.05, both 4C replicates in PDE^−/−^ limbs and FourCseq P > 0.15, both 4C replicates in wild type limbs; Table S1). However, we note that due to the limited sensitivity of 4C, we cannot rule out the formation of transient chromatin interactions or interactions restricted to a subset of cells in the limb bud.

To identify potential compensatory increases in enhancer activity (independent of topology), we used ChIP-seq to profile H3K27ac in E11.5 hindlimbs derived from wild type and PDE^−/−^ mice. Compensatory enhancer activation events may appear as new H3K27ac peaks, or as increases in H3K27ac at active enhancers. We also profiled H3K27me3 in the E11.5 PDE^−/−^ and wild type hindlimb to identify any potential changes to repressed sequences in the PDE^−/−^ background. Genome-wide, we identified 11 H3K27ac and 33 H3K27me3 regions showing statistically significant differential marking between the wild type and PDE^−/−^ samples (DESeq BH *P* < 0.01; Table S6-7). The *Pitx1* gene body itself saw a significant overall loss of acetylation of about 10% in the PDE^−/−^ mice (DESeq BH *P* = 4.33E-05), consistent with the loss of expression we observed. Two other significant changes in H3K27ac signal were found on chromosome 13, though neither was located within the *Pitx1* TAD. These were comprised of a 15% gain and a 10% loss 4.5 Mb and 25 Mb away, respectively, from *Pitx1.* Neither of these sites showed evidence of significant interactions with the *Pitx1* promoter in our 4C data in either wild type or PDE^−/−^ hindlimb. Thus, we found no evidence of new topological interactions or compensatory activity at existing enhancers in the *Pitx1* TAD in E11.5 hindlimbs from PDE^−/−^ mice (Fig 4).

**Figure 4.**
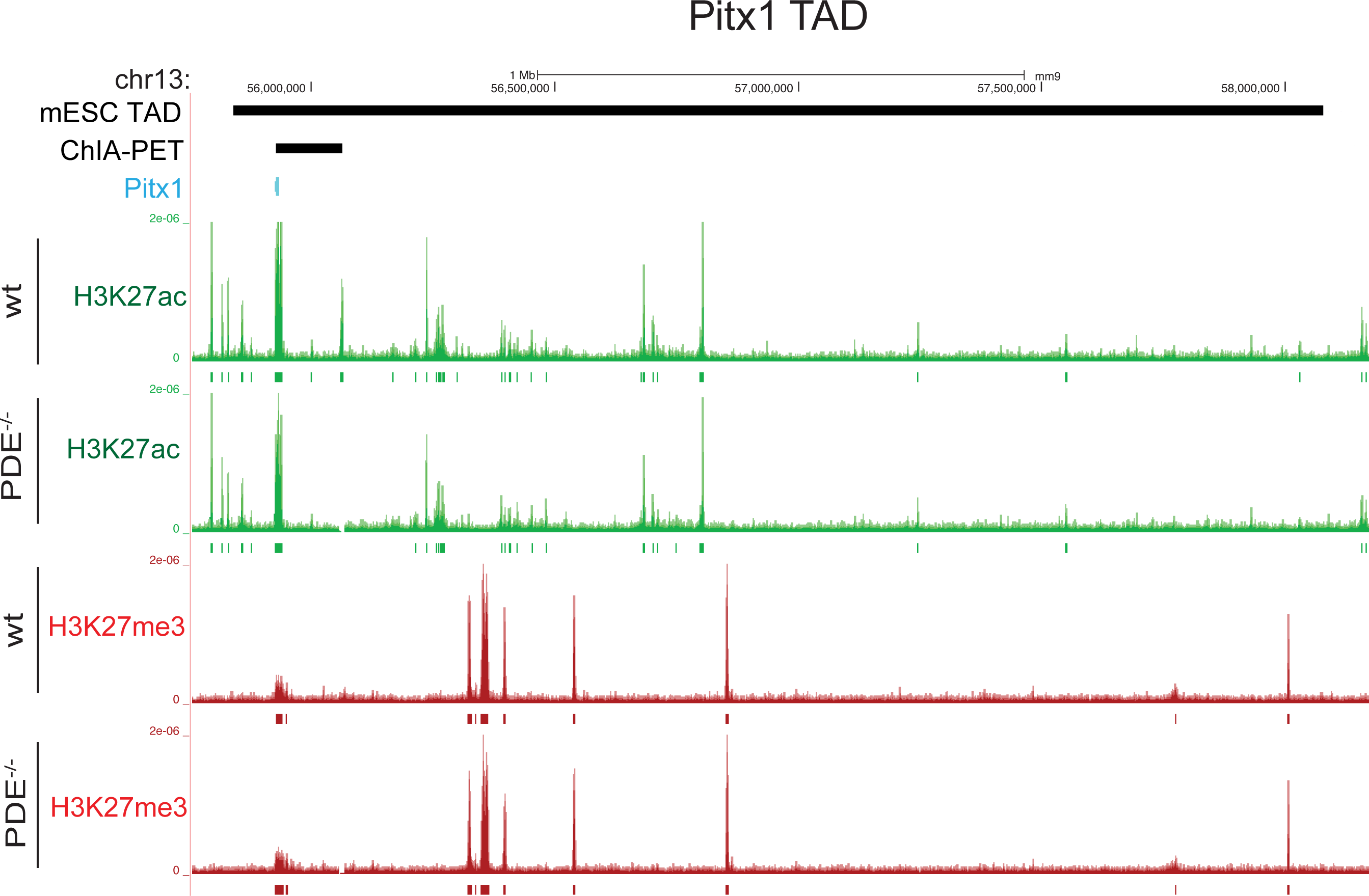
Epigenetic profiles are maintained within the *Pitx1* TAD in wild type and PDE−/− hindlimbs. H3K27ac (green) and H3K27me3 (red) profiles in E11.5 hindlimb from wild type and PDE−/− mice. The TAD identified in Ref × that encompasses *Pitx1* is shown in black. Enriched regions identified for each mark in each tissue are shown below each signal track.

### Morphological analysis of PDE^−/−^ mice

We also evaluated the impact of the *Pitx1* PDE deletion on skeletal morphology. Previous studies have described the complete loss of the ilium and patella in *Pitx1*^−/−^ mice (31). These mice also exhibit a reduced hindlimb stylopod length and a malformed mandible. We examined skeletal morphology in twenty PDE^−/−^ mice at stage E18.5 using Alizarin red/Alcian blue staining. None of these showed any of the gross anatomical defects observed in *Pitx1*^−/−^ mice (Fig. 5, Fig. S5-6). Both the overall morphology and length of the hindlimb stylopod and mandible appeared unchanged in PDE^−/−^ mice.

**Figure 5.**
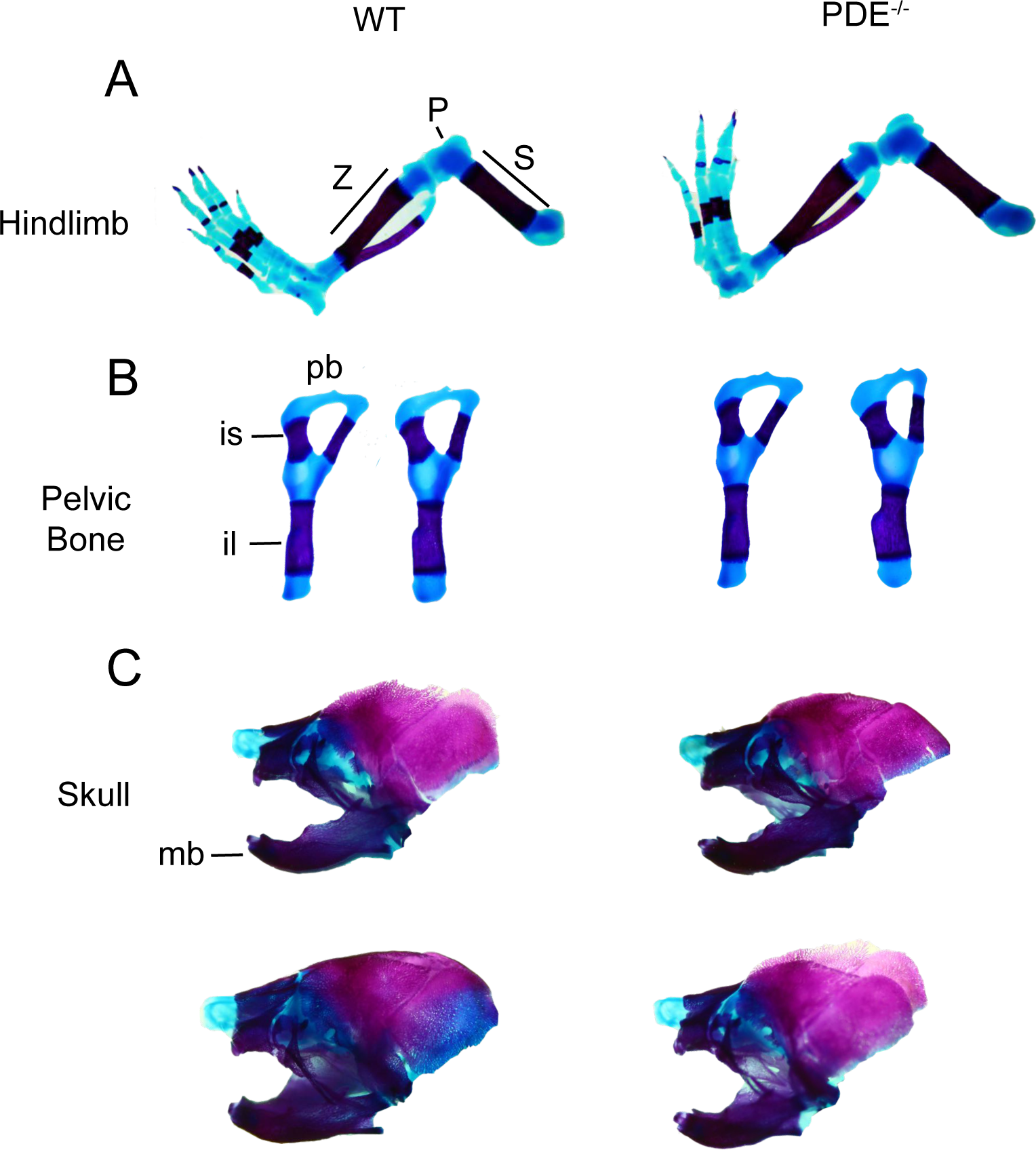
Comparison of skeletal morphology in wild type and PDE−/− E18.5 mouse embryos. Representative alcian blue (cartilage) and alizarin red (bone) skeletal preparations are shown for each genotype. A. Wild type and PDE−/− hindlimbs. S = stylopod; Z = zeugopod; P = patella. B. Pelvic bones from wild type and PDE−/− mice. il = ilium; is = ischium; pb = pubic bone. C. Wild type and PDE−/− skulls. The location of the mandible (mb) is indicated.

To detect potentially subtle reductions of the hindlimb we measured the length of the stylopod and zeugopod in litter-matched E18.5 pups. In our comparisons, we considered the ratio of the length of the stylopod and zeugopod (S/Z) to normalize for any natural variation in overall embryo size. We measured each limb segment both by the length of ossified bone alone as well as the combined length of the bone and associated cartilage. Across multiple litter comparisons the average reduction of the S/Z ratio was under 2%. This reduction was not statistically significant using either metric (ossified bone two-way mixed effect ANOVA F_1,18_ = 2.32, *P* = 0.145; combined bone and cartilage ANOVA F_1,18_ = 0.032, *P* =0.859; Table 1).

**Table 1.** Comparisons of stylopod/zeugodpod length ratios in litter matched E18.5 wild type and PDE−/− hindlimbs. For each litter, the average (S/Z) ratio in PDE−/− hindlimbs was divided by the (S/Z) ratio in wild type hindlimbs.

|  | bone | bone and cartilage |
| --- | --- | --- |
| Litter 1 | 96.47% | 92.00% |
| Litter 2 | 97.45% | 95.72% |
| Litter 3 | 94.91% | 96.27% |
| Litter 4 | 102.25% | 105.97% |
| Litter 5 | 97.29% | 101.81% |
| Litter 6 | 101.13% | 99.25% |
| Average | 98.25% | 98.51% |
| Mann-Whitney U | 0.08364 | 0.41794 |
| ANCOVA | F1,18 = 2.32, P = 0.145 | (F1,18 = 0.032, P = 0.859) |

## Discussion

In this study, we disrupted the predominant topological interaction partner of the *Pitx1* promoter in the hindlimb. Based on previous studies and our 4C results, we predicted the PDE would provide substantial quantitative, spatial or temporal regulatory input to *Pitx1*, and that PDE^−/−^ mice might exhibit profound molecular and morphological phenotypes due to loss of *Pitx1* expression in the hindlimb (10, 30, 31). However, our findings show that the deletion of the PDE and the resulting loss of the major topological interaction of the *Pitx1* promoter does not result in reduced viability or recapitulation of the morphological phenotypes of *Pitx1*^−/−^ mice (31). Although we did not observe major spatial or temporal changes in *Pitx1* expression in PDE^−/−^ mice, our findings do support that the PDE provides quantitative regulatory input to *Pitx1*. The loss of the PDE results in reduced *Pitx1* expression in both the developing hindlimb and mandibular arch, as well as reduced H3K27ac signal at the *Pitx1* promoter. This is not sufficient to substantially alter hindlimb or jaw morphology in our model, indicating the development of these structures is robust to moderate reduction in *Pitx1* dosage. It is possible that PDE^−/−^ mice exhibit hindlimb or jaw defects at very low penetrance, or subtle phenotypes that might be revealed by detailed morphometric analyses that are beyond the scope of our study (19).

Compensatory regulatory interactions that buffer *Pitx1* expression against the loss of the PDE may account for the moderate molecular effects of disrupting the PDE-*Pitx1* interaction. These compensatory interactions may involve recruitment of novel enhancers in the *Pitx1* locus or upregulation of enhancers already in use. However, we detected no evidence of robust novel interactions or enhancer upregulation in hindlimbs of PDE^−/−^ mice. It is possible that transient or weak compensatory interactions may be occurring that our 4C methods are not sensitive enough to detect. However, the absence of a novel robust interaction similar to the PDE-*Pitx1* interaction in PDE^−/−^ mice, coupled with the lack of quantitative changes in H3K27ac marking in the *Pitx1* TAD, suggests that *Pitx1* regulatory architecture is largely unchanged by disruption of the PDE. Moreover, any compensatory mechanisms that may exist are necessarily incomplete, as deletion of the PDE has a clear negative effect on *Pitx1* expression levels.

The PDE may thus provide only a moderate contribution to *Pitx1* regulation, despite its prominent interaction with the *Pitx1* promoter. Other regulatory elements in the *Pitx1* locus, which may be partially redundant with the PDE, must determine the spatial, temporal and quantitative expression of *Pitx1*. Regulatory redundancy is a well-established feature of many developmental genes (4, 5, 22, 23, 25, 26, 41). Our 4C analysis did not reveal other robust, long-range interactions that may be redundant with the *Pitx1*-PDE interaction in the hindlimb. However, H3K27ac profiles in the hindlimb suggest additional enhancers may be located near the *Pitx1* promoter itself. We detected high levels of H3K27ac marking across the *Pitx1* gene body, including intronic marking that may point to proximal enhancers (Fig. 1). Interactions with such proximal elements cannot be distinguished from background in 4C, given their location near the *Pitx1* promoter viewpoint. Distal enhancers within the *Pitx1* TAD may also act redundantly with the PDE (Fig. 4) by interacting with the *Pitx1* promoter at levels below the sensitivity of our 4C methods.

In light of our results, we speculate that the PDE may nevertheless be critical for *Pitx1* expression and normal development in ways that cannot be measured in a laboratory setting. The PDE is highly conserved, suggesting it serves an essential function. Redundant enhancers are thought to be maintained by selection in part because they confer a degree of regulatory robustness that buffers developmental gene expression against perturbation (22, 25, 26). Embryonic development in wild populations is subject to environmental pressures and insults not present in experimental systems. The primary regulatory contribution of the PDE could be to stabilize *Pitx1* expression against variation during development. In the absence of the PDE, *Pitx1* expression may exhibit higher levels of sensitivity to environmental factors or genetic variation, and potentially show much greater changes than we can detect in our model.

Our results also have implications for using topology maps to identify critical regulatory interactions for developmental genes. A primary motivation for generating large-scale maps of topological interactions in the human genome is to discover noncoding variants that contribute to disease by disrupting long-range regulation and perturbing gene expression (12). However, as the PDE illustrates, robust, tissue-specific interactions may not serve to predict enhancer mutations that may have strong effects in model systems. Although it has become commonplace to globally map putative regulatory interactions in multiple biological contexts, we still lack the means to distinguish essential versus redundant interactions *a priori*. This will require large-scale genetic disruption of many interactions, individually and in combination, in order to empirically measure the distribution of effects and characterize the functional diversity of regulatory interactions in the genome.

## Materials and Methods

### Generating PDE^−/−^ conditional knockout alleles using genome editing

All animal work was performed in accordance with approved Yale IACUC protocols. Genetically modified mice were generated at the Yale Genome Editing Center (42). sgRNAs were selected for minimal off-target effects based on a CRISPR Design Tool score of >70 (http://crispr.mit.edu/) and the absence of target sites with 3 or less mismatches on the same chromosome (chr13) as the targeted sequence. Mice were injected with embryo microinjection buffer (10 mM Tris-HCl pH 7.4, 0.25 mM EDTA) containing two guide RNAs each at 50ng/ul, Cas9 RNA at 100ng/ul and two corresponding single stranded DNA donor sequences both at 100ng/ul filtered at 0.22μM (all sequences in Table S8). The Cas9 and sgRNA *in vitro* transcription templates were produced via PCR using a px330 plasmid in which the sgRNAs had been cloned in as described by Zhang et al. as a template (http://www.genome-engineering.org/crispr/wp-content/uploads/2014/05/CRISPR-Reagent-Description-Rev20140509.pdf), placing the sequences under control of the T7 promoter. The resulting DNA products were subjected to *in vitro* transcription using MEGAshortscript T7 Kit (AM1354) for the sgRNAs and mMESSAGE MACHINE^®^ T7 ULTRA Transcription Kit (AM1345) for Cas9. The transcribed RNAs were then purified with the MEGAclear Transcription Clean-Up Kit (AM1908). All primers used are described in Table S8.

Resulting F0 mice were backcrossed to wild type C57BL/6J. To delete the PDE sequence, these mice were then crossed with an actin-cre C57BL/6J mouse line provided by the Yale Genome Editing Center. All mice used in our analysis were from the F3 generation or later.

### 4C analyses in hindlimbs from wild type and PDE^−/−^ mice

100 hindlimbs and 100 forelimbs were dissected from stage E11.5 C57BL/6J embryos. The tissue was crosslinked and processed in accordance with a protocol adapted from Naumova et al. (37) and van de Werken (38) (associated 4C primers listed in Table S8, 4C protocol included in Supplemental Methods). The resulting libraries were prepped and sequenced (1 × 75 bp) on an Illumina Hiseq 2500. For sequencing, the samples were indexed and multiplexed over two lanes. 30% phiX DNA was doped into each sequencing run to mitigate Illumina sequencing artifacts arising due to the low complexity of PCR-amplified 4C libraries.

After sequencing, the 5’ and 3’ primer sequences were removed from the raw reads using cutadapt (43) (v1.4.1) (cutadapt –discard-untrimmed −g $firstprimer −n 10 −m 10 −O 10) and the trimmed reads were then aligned to the mm9 reference genome using bwa (44) (v0.7.10) (bwa mem −t 4). These aligned reads (ranging from 7.5-11 million per replicate) were then used to build a statistical model as described in Klein et al. (36) modified to incorporate aligned sequence from both ends of paired end reads. Raw reads were converted to wiggle tracks using bedtools (45) (v2.17.0) for visualization.

### *RT-qPCR analysis of* Pitx1 *expression*

Hindlimb RNA was collected from five 1:1 litter matched pairs of mice at four different developmental time points: E10.5, E11.5, E12.5, and E13.5 (5 litters from each time point, one wild type and one PDE^−/−^ from each litter, 40 total mice considered). RNA was purified from hindlimb pairs from each individual using the miRNeasy micro (cat#74004) for E10.5 samples and miRNeasy mini (cat#74106) for all others. RNA quality was measured using an Agilent RNA 6000 Pico chip; all samples analyzed had RIN values ≥ 8. We used 100ng of total RNA to prepare cDNA from each sample using the Invitrogen Superscript III reverse transcription kit (cat#18080-051). The resulting cDNA was used in qPCR reaction using Thermo Power SYBR Green mastermix (cat#4367659) (qPCR primers listed in Table S8). All samples were run in sets of triplicates reporting the Ct values of *Pitx1* and *Hprt1*, which we used as an internal reference. Reductions in gene expression were calculated by comparing ^ΔΔ^Ct values of the PDE samples to wild type.

### Global transcriptome analyses in hindlimbs and mandibular arch from wild type and PDE^−/−^ mice

RNA was collected from hindlimb bud pairs of four litter-matched E11.5 mice (2 wild type, 2 PDE^−/−^) generated from a PDE^+/−^ × PDE^+/−^ cross. The mandibular arch analysis RNA was derived from three 1 to 1 pairs of litter matched E11.5 mice (3 litters, one wild type and one PDE^−/−^ from each). All RNA was purified using the Qiagen miRNeasy RNA extraction kit (74106). Prior to sequencing, RNA quality was analyzed on an Agilent 2100 RNA pico chip; all samples submitted for sequencing had RIN values ≥ 8. RNA was library prepped and sequenced using standard Illumina protocols (2 × 75bp) on an Illumina HiSeq 2500 (mandibular arch) and Illumina HiSeq 4000 (hindlimb). Raw reads were aligned to the mm9 transcriptome (GRCm38.p4) using tophat2 (46) (v2.0.9) and Gencode vM.7. Statistical analysis of expression changes was conducted using DESeq. (39) using default settings and an FDR of 0.1.

### *Analysis of* Pitx1 *expression using* in situ hybridization

*In situ* hybridization was performed at E11.5 as previously described (47), using a *Pitx1* probe provided by Jacques Drouin (48).

### Global profiling of H3K27ac and H3K27me3 in hindlimbs from wild type and PDE ^−/−^ mice

We performed ChIP-seq for each epigenetic mark using chromatin derived from 180 wild type and 180 PDE^−/−^ hindlimbs (30 per replicate, 3 replicates per mark) from over 10 litters. Hindlimb tissue was crosslinked, pooled, and sonicated. The pools of each genotype were then split into three ChIP replicates for each epigenetic mark and processed independently for the remainder of the protocol. ChIP-seq of H3K27ac (2ug/ChIP, Active Motif 39155) and H3K27me3 (5ug/ChIP, Active Motif ab4729) was performed as described in Cotney *et al*. 2012 (36). Samples were sequenced (2 × 75b) on an Illumina 2500. To control for batch effects all H3K27ac samples were multiplexed and sequenced on a single lane. The same method was used for H3K27me3 samples. Raw reads were aligned to the mm9 reference genome using bowtie2 (v2.2.3) and peaks were called using MACS (49) (v 1.4.2).

For quantitative analysis, H3K27ac or H3K27me3 peaks that were reproducible in all three replicates for each genotype were merged. Peaks were considered reproducible if they overlapped by at least 1 bp (identified with the default settings of the intersectBed command in BEDTools). The relative enrichment of H3K27ac and H3K27me3 in these peaks was quantitatively compared using DESeq2 and an FDR of 0.1.

### Analyses of skeletal morphology in E18.5 wild type and PDE^−/−^ mice

E18.5 skeletons were stained with Alcian blue and Alizarin red as previously described (47). Litter matched mice from six litters were considered represented by 30 wild type and 24 PDE^−/−^ independent hindlimbs. Bone and cartilage lengths of the stylopod and zeugopod portions of the hindlimb were photographed under a stereo microscope (Leica S6 D) and the segment length were measured independently and blinded to genotype by two individuals using Photoshop CC (2017.0.0).

### ANCOVA and ANOVA analyses

To dissect the main effects of the factors contributing to the observed differences in RT-qPCR and limb growth measurements, we analyzed these datasets in comprehensive modeling frameworks. For the RT-qPCR dataset, we employed a three-way mixed-effect ANCOVA with ^ΔΔ^CT measurements as the response variable, ‘genotype’ as a two-level fixed effect factor, ‘embryonic age’ [in days] as a covariate, and ‘litter’ as a random effect factor that is nested within ‘embryonic age.’ For the limb measurements, we used a two-way mixed-effect ANOVA with stylopod-to-zygopod ratio (both accounting for bone only and for bone and cartilage together) as the response variable, ‘genotype’ as a two-level fixed effect factor, and ‘litter’ as a random effect factor. This way we were able to isolate the effects of ‘genotype,’ our factor of interest, from the potential effects of other contributing factors.

All associated raw sequence data has been submitted to the SRA (Submission ID: SUB2725331, BioProject ID: PRJNA388175).

## Acknowledgments

Our thanks to Jacques Drouin for providing the *Pitx1 in situ* probe. We also thank members of the Noonan laboratory for insightful comments on the manuscript. This work was supported by a grant from the National Institute of General Medical Sciences (GM094780) to J.P.N. and F32 GM106628 (to D.E.), a grant from the Deutsche Forschungsgemeinschaft (UE 194/1-1) to S.U. R.S. was supported by National Institutes of Health training grants T32 GM007223 and T32 HD007149. This work was supported by the HPC facilities operated by, and the staffs of, the Yale Center for Research Computing and the Yale Center for Genome Analysis.

